# CRISPR Activation/Inhibition Experiments Reveal that Expression of Intronic MicroRNA *miR-335* Depends on the Promoter Activity of its Host Gene *Mest*

**DOI:** 10.1101/2021.09.15.458166

**Authors:** Mathilde Courtes, Céline Lemmers, Anne Le Digarcher, Ilda Coku, Arnaud Monteil, Charles Hong, Annie Varrault, Tristan Bouschet

**Affiliations:** Institut de Génomique Fonctionnelle, Université de Montpellier, CNRS, INSERM, Montpellier, France; Plateforme de Vectorologie de Montpellier (PVM), BioCampus Montpellier, Université de Montpellier, CNRS, INSERM, Montpellier, France; Vanderbilt University School of Medicine Nashville, Nashville, USA

**Keywords:** microRNA, CRISPR, CRISPRa, CRISPRi, pluripotent stem cells, mouse, Mest, miR-335

## Abstract

MicroRNAs are small non-coding RNAs that act as rheostats to modulate gene expression during development, physiology, and disease. Approximately half of mammalian microRNAs are intronic. It is unknown whether intronic miRNA transcription depends on their host gene or a microRNA-specific promoter. Here, we show that CRISPR inhibition of host gene *Mest* downregulated hosted *miR-335* in mouse embryonic stem cells and brain organoids. Reciprocally, CRISPR transactivation of *Mest* upregulated *miR-335*. By contrast, activation of *miR-335* predicted promoter had no effect. Thus, intronic *miR-335* expression depends on the promoter activity of its host gene. This approach could serve to map microRNA promoters.

## INTRODUCTION

microRNAs (miRNAs) are short non-coding RNAs that play a central role in regulating gene expression in plants and animals (Bartel 2018; Jones-Rhoades et al. 2006). miRNAs impact on development and physiology, and are dysregulated in diseases, including cancer (DeVeale et al. 2021; Schanen and Li 2011; Xue et al. 2021). Stringent gene annotations suggest that there are ∼500 miRNAs in mice (Chiang et al. 2010) and humans (Fromm et al. 2015). It is estimated that approximately half of mammalian miRNAs are intronic (Meunier et al. 2013; Rodriguez et al. 2004; Hinske et al. 2014).

miRNAs biogenesis sequentially involves transcription, cleavage of the miRNA hairpin precursor out of the primary transcript, transport of intermediate forms, and loading of the mature miRNA into the RNA-induced silencing complex (Bartel 2018; Westholm and Lai 2011; Ha and Kim 2014). The mechanisms that regulate miRNAs transcription, a key factor of miRNA abundance and tissue-specific expression, are not well defined, in particular for intronic miRNAs. Intronic miRNAs were first observed as frequently co-regulated with their host genes (Baskerville and Bartel 2005; He et al. 2012; Liang et al. 2007; Rodriguez et al. 2004; Seitz et al. 2004), suggesting that their transcription depends on the promoter activity of the host gene. By contrast, recent work suggests that most intronic miRNAs are not co-regulated with their host genes, which is supported by the fact that they have independent transcription start sites (Steiman-Shimony et al. 2018). Many additional studies have tried to map miRNA promoters using bioinformatics tools (Chen et al. 2019). For instance using chromatin modifications (Ozsolak et al., 2008) or deepCAGE data (Marsico et al., 2013) it was estimated that ∼30% of intronic miRNAs have independent promoters. To our knowledge, whether the transcription of an intronic miRNA depends on the promoter activity of the host gene or a miR-specific promoter has not been tested experimentally.

*Mest* (Mesoderm-specific transcript) is a protein-coding gene that hosts *miR-335* in one of its introns. *Mest* and *miR-335* are highly conserved during evolution and frequently co-regulated (Hiramuki et al. 2015; Liang et al. 2007; Ronchetti et al. 2008; Tomé et al. 2011; Yang et al. 2014). This suggests that *Mest* and *miR-335* are controlled via common regulatory sequences, possibly *Mest* promoter. In addition, based on a luciferase assay, *miR-335* was proposed to have an independent promoter located in a *Mest* intron (Zhu et al. 2014).

Here, to get insights into the mechanisms of transcription of an intronic miRNA, we have applied CRISPR/Cas9 -Clustered regularly interspaced short palindromic repeats (CRISPR) and CRISPR-associated protein 9 (Cas9)-technologies to *Mest* and intronic *miR-335*. More specifically, we have used CRISPR activation (CRISPRa) and CRISPR inhibition (CRISPRi) where a cleavage defective Cas9 (dCas9) is fused to either activators or repressors of transcription (Konermann et al. 2015; Yeo et al. 2018; Gilbert et al. 2013) and directed these complexes to the endogenous promoter sequences of *Mest* or to the predicted promoter of *miR-335*.

## RESULTS & DISCUSSION

### CRISPRi of host gene *Mest* suppresses the expression of hosted *miR-335* in embryonic stem cells

*miR-335* is located in an intron of the protein-coding gene *Mest* (Fig. 1A, B) and is transcribed from the same DNA strand as its host gene, a common feature of intronic miRNAs (Hinske et al. 2014). *Mest* has one distal promoter (D) and one proximal promoter (P). *Mest* is highly expressed in mouse embryonic stem cells (mESCs) and *Mest* transcripts originate predominantly from the proximal promoter (P) (Fig. 1A, generated from previously published RNA-seq experiments (Bouschet et al. 2017)). Furthermore, miR-335-3p was reported to be expressed in mESCs (Kingston and Bartel 2019).

**Figure 1.**
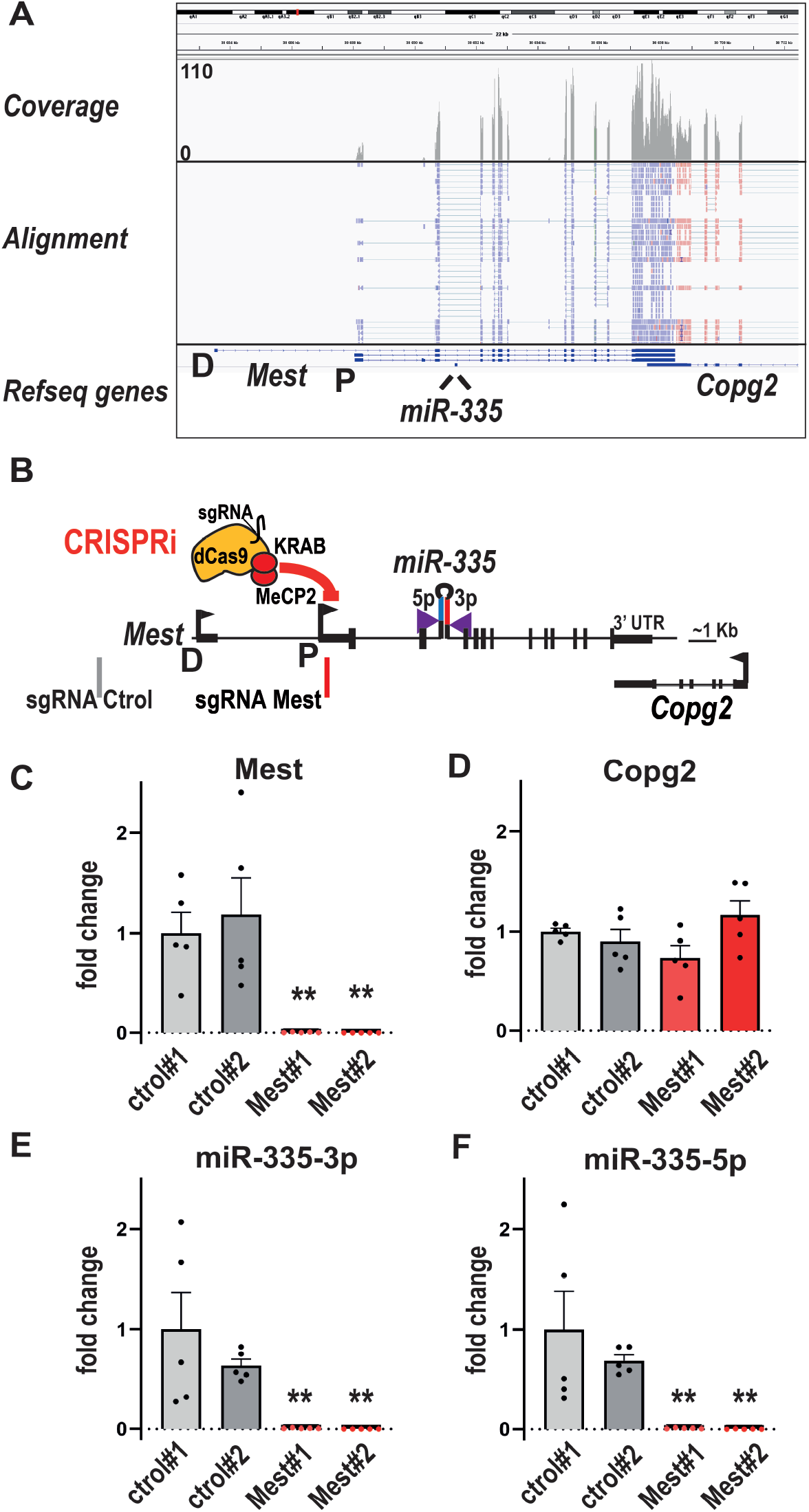
CRISPRi on *Mest* suppresses the expression of hosted *miR-335* in embryonic stem cells. (A) Transcription originates from the proximal promoter of *Mest* (P) in mouse embryonic stem cells. Integrative Genomics Viewer tracks showing coverage plot and alignment of RNA-seq reads for mouse embryonic stem cells. Reads for *Mest* (blue) are transcribed from the plus strand, while reads from *Copg2* (pink) are transcribed from the minus strand. Chromosomal coordinates and gene annotation are from the RefSeq mm9 build. D: *Mest* distal promoter; P: *Mest* proximal promoter. (B) Schematic of mouse *Mest* gene with the CRISPRi module (dCas9-KRAB-MeCP2) targeting the proximal promoter P of *Mest*. (C, D) Repression of *Mest* promoter downregulates *Mest* (C) but does not affect the expression of neighboring *Copg2* (D). RNAs were quantified in two CRISPRi ESC clones expressing the control sgRNA (grey) and two CRISPRi ESC clones expressing *Mest* sgRNA (red). Data are mean ± sem of five independent experiments and expressed as fold change over control clone #1. **:p<0.01 (Mann-Whitney test). (E, F) Influence of repressing *Mest* promoter on miR-335-3p and miR-335-5p levels. Data are mean ± sem of five independent experiments and expressed as fold change over control clone #1. **:p<0.01 (Mann-Whitney test).

We reasoned that if *miR-335* expression depends on the activity of *Mest* promoters, then repressing transcription at *Mest* promoters in mESCs with CRISPRi should decrease miR-335 transcripts. Using Hyper-piggyBac transposase (Yusa et al., 2011), we first generated a CRISPRi mESC line that stably expressed dCas9 fused to the repressors of transcription KRAB and MeCP2. dCas9-KRAB-MeCP2 was previously shown to efficiently repress a vast panel of genes in HEK293T cells (Yeo et al. 2018). CRISPRi mESCs (characterized in Supplemental Fig. S1) were transduced with lentiviruses that express either a control sgRNA (no match in the mouse genome) or a sgRNA targeting either the distal or the proximal promoter of *Mest* (Supplemental Fig. S2A). sgRNAs targeting *Mest* proximal promoter P downregulated *Mest* while targeting distal promoter D had no obvious effect (Supplemental Fig. S2B). Thus, as expected, CRISPRi was efficient only when targeting the active *Mest* promoter. Levels of the neighboring gene *Copg2* were unaffected (Supplemental Fig. S2C).

We then selected two CRISPRi mESC clones expressing the control sgRNA and two clones expressing the sgRNA *Mest* P2 for further analyses (Fig. 1B). There was a >100 fold-downregulation of *Mest* in CRISPRi *Mest* clones compared to CRISPRi control clones (Fig. 1C). By contrast, *Copg2* expression was unaffected (Fig. 1D). We next measured the levels of miR-335-3p and miR-335-5p, the final products of miR-335 biogenesis, by gene-specific RT followed by qPCR with Taqman probes. In CRISPRi Mest clones, miR-335-3p and miR-335-5p levels were reduced to less than 1% of levels measured in CRISPRi control clones (Fig. 1E, F), a massive downregulation that paralleled well that of *Mest* (Fig. 1C). Thus, the transcriptional activity of *Mest* proximal promoter is required for the expression of intronic *miR-335* in mESCs.

### *Mest* promoter activity is required for *miR-335* expression in brain organoids

Next, we determined whether *miR-335* expression dependency on *Mest* promoter persists upon differentiation of mESCs into brain organoids. Brain organoids were generated from mESCs according to a published protocol (Eiraku et al. 2008) with slight modifications (see *Materials and Methods*). RNA-seq experiments on these brain organoids show enrichment in Gene Ontology Terms such as ‘forebrain generation of neurons’ after eight days of differentiation and ‘telencephalon development’ and ‘action potential’ after 15 days of differentiation (Bouschet and co-workers, unpublished).

As expected, brain organoids contained neural progenitors of dorsal identity (NESTIN+PAX6+ cells) after 8 days of differentiation, and neurons (TUBB3+ cells), including some neurons that expressed the cortical marker TBR1 after 15 days of differentiation (Fig. 2A).

**Figure 2.**
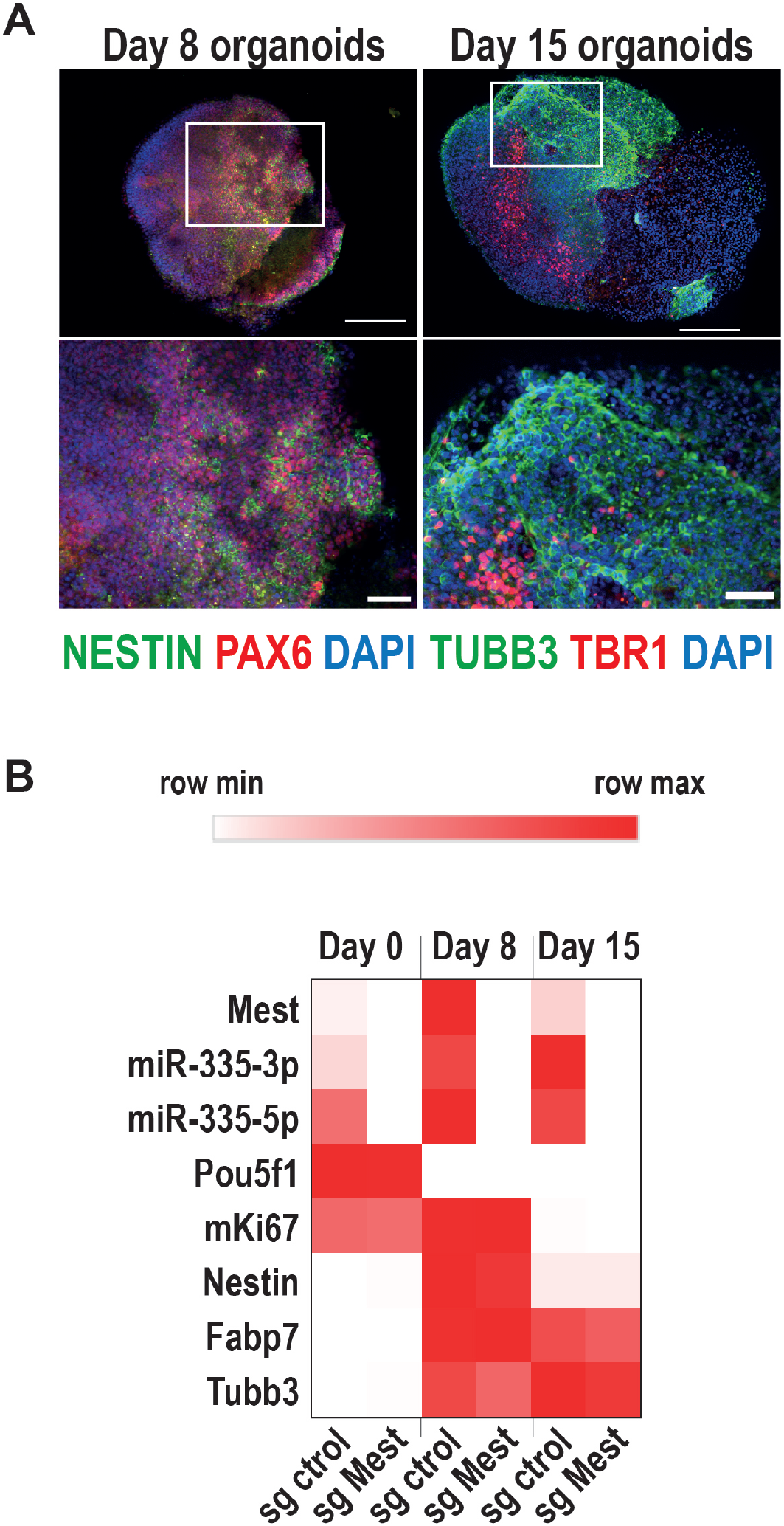
*miR-335* expression depends on *Mest* promoter activity in brain organoids. (A) Immunofluorescence staining on brain organoids derived from mESCs using antibodies for brain primordium markers NESTIN/PAX6 (middle panels) and TUBB3/TBR1 (right panels) after eight and 15 days of differentiation. The top panels show entire organoids. The bottom panels are zoom-in insets of an area in organoids. Scale bars: 200 µm for organoids (top panels) and 50 µm for insets (bottom panels). (B) Time course of expression of *Mest* and *miR-335* mature products during the development of brain organoids from CRISPRi ESCs stably expressing either control sgRNA or *Mest* sgRNA. Heatmap shows the mean of four independent experiments performed on two CRISPRa sgRNA control and two CRISPRa sgRNA ESC clones. Heatmap was built using Morpheus. https://software.broadinstitute.org/morpheus/

*Mest* and *miR-335* transcripts were upregulated during the generation of brain organoids from CRISPRi mESCs expressing the control sgRNA (Fig. 2B). By contrast, *Mest* RNA was barely detectable in CRISPRi organoids expressing the *Mest* sgRNA (Fig. 2B), showing that *Mest* promoter remains repressed in differentiated cells. Importantly, *miR-335* mature products were also barely detectable in these organoids (Fig. 2B). Thus, the activity of *Mest* promoter is required for *miR-335* expression in both undifferentiated mESCs and their neural progeny.

### CRISPRa on *Mest* increases the expression of hosted *miR-335*

We next tested whether transactivating *Mest* promoter is sufficient to increase *miR-335* levels and therefore mirrors CRISPRi loss of function experiments. mESCs stably expressing the CRISPRa Synergistic Activation Mediator (SAM) module (Bonev et al. 2017) -composed of three transactivators (Konermann et al. 2015)- were transduced with lentiviruses expressing either a control sgRNA or a sgRNA targeting Mest (D) or (P) promoter (Supplemental Fig. S3A) -as described for CRISPRi-.

Transactivating *Mest* distal promoter efficiently increased *Mest* transcripts (Supplemental Fig. S3B). By contrast, transactivating *Mest* proximal promoter with sgRNAs P1 and P2 had no major effect on *Mest* transcript level (Supplemental Fig. S3B), likely because this promoter is already very active in ESCs (Fig. 1A). The level of *Copg2* was not altered by any of the three *Mest* sgRNAs (Supplemental Fig. S3C).

We selected for further analysis two CRISPRa control clones and two CRISPRa *Mest* clones (expressing the D sgRNA, Fig. 3A). On average, there was a 3.2 fold increase in *Mest* transcript in CRISPRa *Mest* clones compared to clones expressing the control sgRNA (Fig. 3B). As for CRISPRi, *Copg2* expression was unaffected (Fig. 3C). Strikingly, the levels of both miR-335-3p and miR-335-5p also increased by a ∼3-fold (Fig. 3D, E). Thus, activating the distal promoter of *Mest* with CRISPRa/SAM is sufficient to increase hosted *miR-335* levels in mESCs.

**Figure 3.**
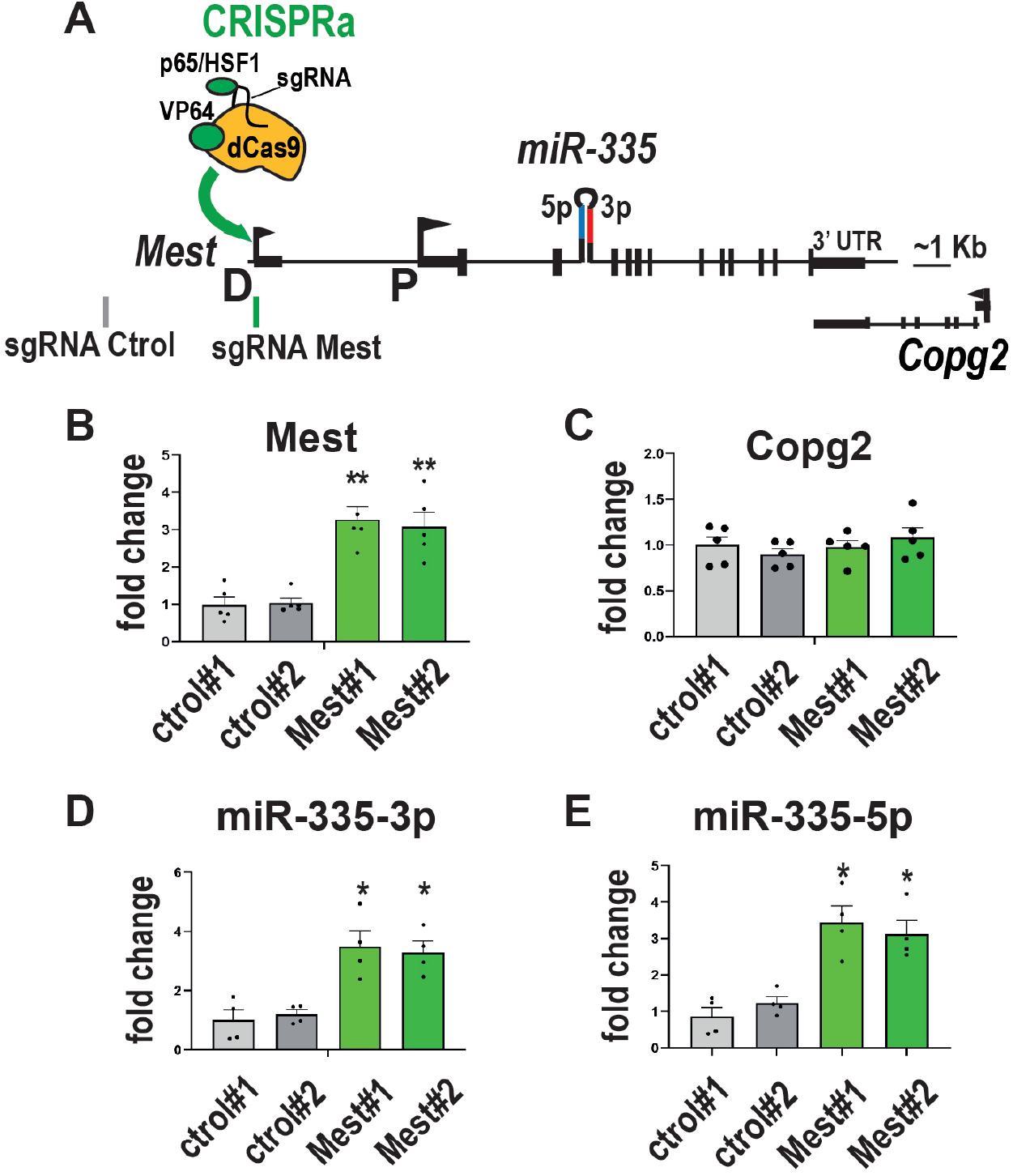
CRISPRa on *Mest* increases the expression of hosted *miR-335* in embryonic stem cells. (A) Schematic of mouse *Mest* gene structure with the CRISPRa SAM - synergistic activation mediator-module targeting the distal promoter D of *Mest*. (B, C) Transactivation of *Mest* promoter upregulates *Mest* (B) but does not affect neighboring *Copg2* expression (C). Data are mean ± sem of five independent experiments performed on two CRISPRa sgRNA control (grey) and two CRISPRa sgRNA *Mest* clones (green) and expressed as fold change over control clone#1.**:p<0.01 (Mann-Whitney test). (D; E) Transactivation of *Mest* promoter increases miR-335-3p (D) and miR-335-5p (E) levels. Data are mean ± sem of four independent experiments and expressed as fold change over control clone#1. *:p<0.05 (Mann-Whitney test).

### CRISPRa on *miR-335* putative promoter does not affect *miR-335* levels

A previous study, based on luciferase assays performed in HEK293T cells, suggests that the sequence upstream of *miR-335* (situated in a *Mest* intron) has some promoter activity (Zhu et al. 2014). Thus, we next tested whether we could upregulate *miR-335* by directing SAM to this genomic region.

Because SAM efficiency correlates with baseline expression levels of the targeted gene – the fold of upregulation is inversely correlated with basal transcript level- (Konermann et al. 2015), and to maximize the chance to increase miR-335, SAM experiments were performed on cells with lower baseline levels of *miR-335* than mESCs. We observed that miR-335-3p and miR-335-5p levels were respectively 13 and 47 times lower in MEFs compared to mESCs (Fig. 4A, B). Mest expression was also ∼60 times less expressed in MEFs than in mESCs (Fig. 4C), adding further support for the coregulation of *Mest* and *miR-335*.

**Figure 4.**
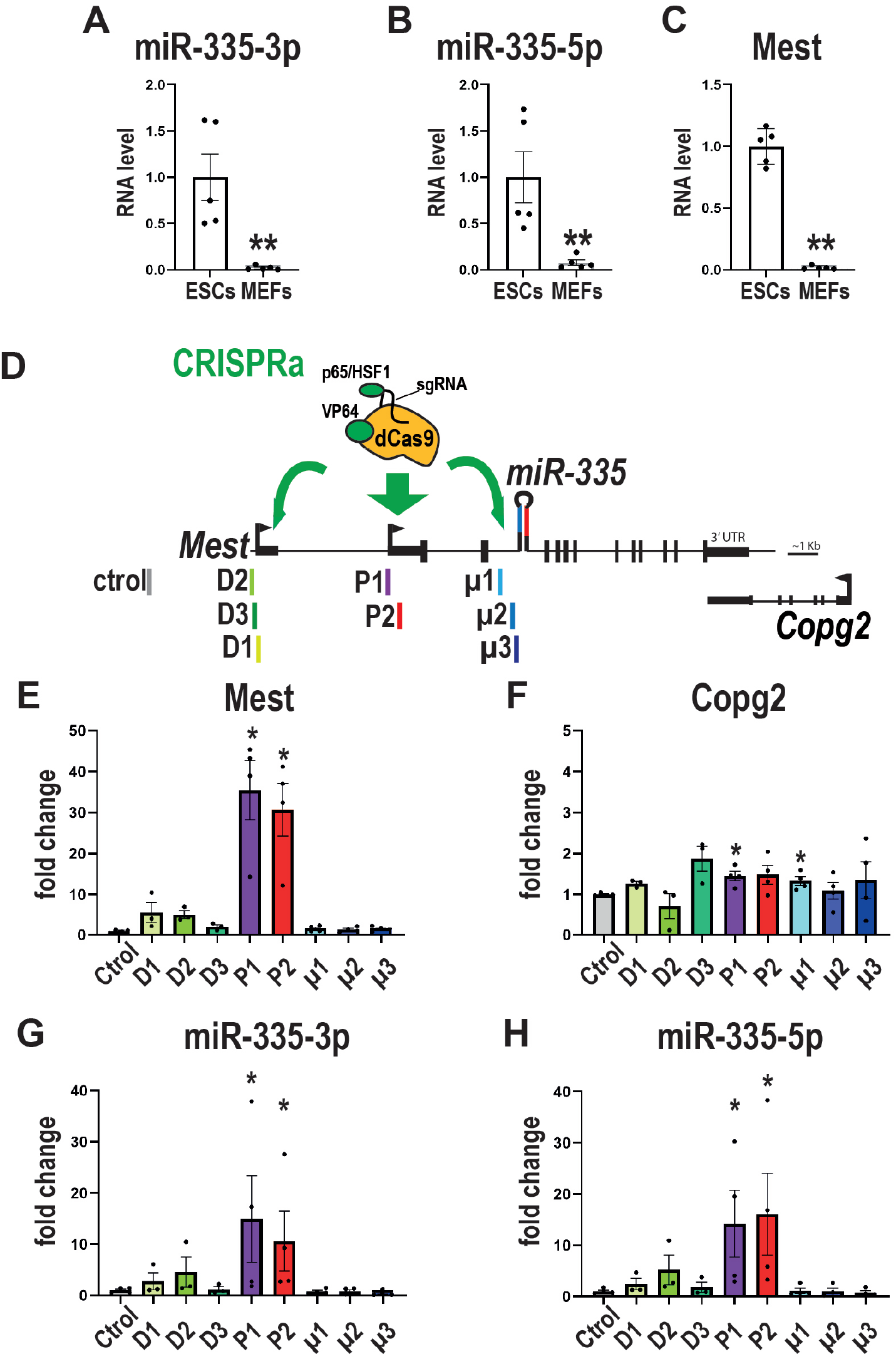
CRISPRa on *miR-335* putative promoter does not affect *miR-335* levels in MEFs. (A-C) Endogenous expression of *miR-335* products and *Mest* is weaker in SAM MEFs than in SAM ESCs. Data are mean ± sem of qPCR experiments performed on five MEF and five ESC samples and normalized to the average value obtained on ESCs.**:p<0.01 (Mann-Whitney test). (D) Structure of mouse *Mest* gene with SAM targeting either the distal promoter of *Mest* D (D1-D2 sgRNAs), the proximal promoter of *Mest* P (P1-P3 sgRNAs), or the putative promoter of *miR-335* (µ1-µ3 sgRNAs). (E-H) Levels of expression of *Mest* (E), *Copg2* (F), *miR-335-3p* (G), and *miR-335-5p* (H) were measured after transactivation of either *Mest* D or P promoters or *miR-335* putative promoter. SAM MEFs were transfected with plasmids expressing sgRNAs targeting *Mest* D (sgRNAs D1, D2, and D3), *Mest* P (P1 and P2), or the putative promoter of miR-335 (µ1, µ2, and µ3). Data are mean ± sem of three to four independent experiments and expressed as fold change over sgRNA control taken as 1. *: p<0.05 in Mann-Whitney test (comparison with sgRNA control values). None of the sgRNAs that direct the CRISPRa/SAM machinery towards the *miR-335* putative promoter (µ1, µ2, and µ3) altered *miR-335* levels.

We designed three sgRNAs (µ1, µ2, and µ3) in the putative *miR-335* promoter –a region named pro2 in (Zhu et al. 2014)- and compared their efficiency in upregulating *miR-335* to sgRNAs that target *Mest* promoters (Fig. 4E). sgRNAs P1 and P2 (which target *Mest* P promoter) strongly upregulated *Mest* (Fig. 4E) but also *miR-335* mature products in SAM MEFs (Fig. 4G, H). The upregulation of *Mest* was much higher in MEFs than in mESCs, as expected from their relative *Mest* baseline levels (see Fig. 4C). By contrast, the three sgRNAs that target the putative promoter of *miR-335* (µ1, µ2, and µ3) did not affect miR-335-3p nor miR-335-5p levels (Fig. 4G, H). Thus, this genomic sequence likely does not regulate miR-335 expression in MEFs. We cannot rule out that *miR-335* has an independent promoter located in another region. In this context, prediction of *miR-335* promoter location using DeepCAGE data (Marsico et al. 2013) suggests that there could be several *miR-335* promoters depending on the tissue. According to Marsico and coworkers, the most probable *miR-335* promoters are *Mest* (D) and (P) promoters - what we confirmed experimentally here-, and less probably, a third region situated in another intron of *Mest*.

Data obtained in MEFs also revealed that transactivating *Mest* (P) promoter resulted in a strong increase in *Mest* and *miR-335* while transactivating (D) promoter had moderate effects. This contrasts with results obtained in ESCs where the most potent sgRNAs were those targeting the (D) promoter (Fig. 3 and Supplemental Fig. S3). Taken together, these data suggest that transcriptional activation of one or the other *Mest* promoter, depending on the cell type, is sufficient to increase the levels of intronic *miR-335*. This also supports the existence of primary transcripts, originating either at (D) or (P) promoters, that contain both *Mest* and *miR-335* precursors.

To conclude, CRISPRa and CRISPRi experiments on *Mest* and *miR-335* in mouse cells reveal that transcription of an intronic miRNA is regulated by the promoter of its host gene. Previous works propose that evolutionarily conserved intronic miRNAs, such as *miR-335*, are more frequently co-expressed with host genes than recently appeared intronic miRNAs (He et al. 2012; Steiman-Shimony et al. 2018). This suggests that the transcription of conserved intronic miRNAs depends on the host promoter while recently appeared intronic miRNAs tend to have independent promoters. To test these predictions our CRISPRa/i approach could be used to map miRNA promoters on a genome-wide scale.

## MATERIALS AND METHODS

### Cell culture

E14Tg2a mouse ESCs and their CRISPRa and CRISPRi derivatives were cultivated on gelatine coated dishes and maintained pluripotent in Serum/Lif media as described (Varrault et al. 2018). Organoids were generated in 96-well (U-bottom) Ultra-Low Attachment plates (Sumitomo) by seeding 3000 ESCs in corticogenesis medium 1: DMEM/F-12/GlutaMAX supplemented with 10% KSR, 0.1 mM of non-essential amino acids, 1 mM of sodium pyruvate, 50U/ml penicillin/streptomycin, 0.1 mM of 2-mercaptoethanol (Sigma), 1 µM DMH1-HCl (in house synthesized, Vanderbilt University) and 240 nM IWP-2 (Tocris). On day 8 of differentiation, organoids were transferred to bacterial plates (Greiner) in corticogenesis medium 2: DMEM/F-12/GlutaMAX supplemented with N2 and B27 (without vitamin A) supplements, 500 µg/ml of BSA, 0.1 mM of non-essential amino acids, 1 mM of sodium pyruvate, 0.1 mM of 2-mercaptoethanol, and 50U/ml penicillin/streptomycin. Immortalized CRISPRa (SAM) MEFs (gift from Giacomo Cavalli’s lab, unpublished) were cultivated in DMEM supplemented with 10% FBS and 50U/ml penicillin/streptomycin. All media components were from Life Technologies unless otherwise stated. Cell lines were routinely tested for the absence of mycoplasma (Mycoalert, Lonza).

### Generation of constructs expressing sgRNAs

sgRNA sequences targeting *Mest* promoters were designed using CRISPick https://portals.broadinstitute.org/gppx/crispick/public (formerly GPP sgRNA Design tool) or manually. sgRNAs that target the putative miR-335 promoter (mm10_dna range=chr6_30740830-30741300) were designed using CHOPCHOP (Labun et al. 2019). Pairs of oligonucleotides (Eurofins) were annealed and subcloned into either sgRNA(MS2) cloning backbone (Addgene Plasmid #61424) or Lenti sgRNA(MS2)_zeo backbone (Konermann et al., 2015) (Addgene plasmid # 61427) that were previously digested with either BbsI or BsmBI (NEB), respectively, and purified on a Chromaspin column (Clontech). All constructs were verified by Sanger sequencing (Genewiz). sgRNA sequences are listed in Supplemental Table S1.

### Lentiviruses production

Lentiviruses were prepared as described elsewhere (Lin et al. 2002). Briefly, lentiviral transfer vectors were co-transfected with the HIV packaging plasmid psPAX2 and the plasmid pMD2G (coding for the vesicular stomatitis virus envelope glycoprotein G), in HEK-293T cells by the calcium phosphate method. Supernatants were collected at day 2 post-transfection and concentrated on sucrose by ultracentrifugation at 95 528g for 1.5 h at 4°C.

### Generation of CRISPRi ESC lines using PiggyBac Transposition

E14Tg2a mouse ESCs were co-transfected with pCMV-HA-HyperpiggyBase (Yusa et al., 2011) and pB-dCas9-KRAB-MecP2 (Yeo et al. 2018) (Addgene plasmid # 110824) using a Neon transfection system (Life Technologies). Forty-eight hours post-transfection, cells were selected using Blastidicin (15 µg/ml, SIGMA). Stable pB-dCas9-KRAB-MeCP2 ESCs (CRISPRi ESCs) were then transduced with lentiviruses expressing the following sgRNAs: control, *Mest* distal promoter, *Mest* proximal promoter#1, or *Mest* proximal promoter#2. Seventy-two hours post-infection, cells were selected using hygromycin (1 mg/ml, Life Technologies), and clones were picked and expanded in ESC media.

### Generation of SAM CRISPRa ESC lines targeting *Mest* promoters

E14Tg2a ESCs stably expressing the SAM system (Bonev et al. 2017) –SAM ESCs-were transfected with Lenti sgRNA(MS2)_zeo plasmids expressing the following sgRNAs: control, *Mest* distal promoter, *Mest* proximal promoter#1, or *Mest* proximal promoter#2. ESCs were selected using Zeocin (250 µg/ml, Life Technologies) and clones were picked and expanded.

### Transient transfection of SAM MEFs

80 000 MEFs stably expressing the SAM system (SAM MEFs) were transfected using Lipofectamine 2000 with 300 ng of sgRNA(MS2) plasmid expressing either one control sgRNA, one *Mest* distal promoter sgRNA (out of 3 different sgRNAs), one *Mest* proximal promoter sgRNA (out of 2 different sgRNAs), or one *miR-335*-putative promoter sgRNA (out of 3 different sgRNAs). Forty-eight hours later, RNAs were harvested.

### RNA extraction and RT-qPCR

Total RNAs were extracted using quick-RNA miniprep kits (Zymo) and quantified on a Nanodrop. RNAs were retro-transcribed with N6 primers and M-MuLV retro-transcriptase (RT). qPCR was performed using validated primers and SYBR Green Mix (Roche) in 384-well plates on a LightCycler480 device (Roche) as described in (Varrault et al. 2018). The level of expression of each gene was normalized to the average expression levels of three housekeeping genes selected with geNorm (Vandesompele et al. 2002): *Gapdh, Tbp*, and *Mrpl32* for ESCs and *Gapdh, Tbp* and *Gusb* for MEFs. qPCR primer sequences are listed in Supplemental Table S2.

miRNAs were retro-transcribed with gene-specific primers and multiscribe RT (Life Technologies). Their levels of expression were measured with TaqMan probes (miRNA Taqman assays # 000546 for miR-335-5p, and # 002185 for miR-335-3p). and normalized to that of U6 snoRNA (assay # 001973) (ThermoFisher). We found that U6 was stably expressed across samples (not shown).

### Visualization of RNA-seq experiment

RNA-seq reads from mESCs -GSE75486 (Bouschet et al. 2017)- were visualized using Integrative Genomics Viewer (Robinson et al. 2011) -version: 2.8.13-.

### Immunofluorescence

Immunofluorescence experiments were performed as described (Varrault et al. 2018) using antibodies directed against (species; provider; catalog number): CAS9 (mouse; Cell signalling; #14697); NANOG (mouse; BD Pharmingen; #560259); NESTIN (mouse; Santa Cruz; sc-33677); PAX6 (mouse; Covance; PRB-278P); POU5F1 (rabbit; Cell signalling; #2840); TBR1 (rabbit, Cell signalling; #49661); TUBB3 (mouse; Covance; MMS-435P). Secondary antibodies were anti-mouse or anti-rabbit coupled to Alexa Fluor® 488 or Cy3 (Jackson Immunoresearch Laboratories). Nuclei were labeled with DAPI and slides were mounted with mowiol and observed under a fluorescence microscope (ImagerZ1, Zeiss). Images of organoids were obtained by tiling and stitching, and insets were taken using the apotome mode.

### Statistical analysis

Statistical analysis was carried out using GraphPad Prism Version 8 (GraphPad Software, San Diego, USA). Mann-Whitney test was used for comparing differences between two groups. p values < 0.05 were considered statistically significant.

## ACKNOWLEDGMENTS

We thank Laurent Journot for helpful discussions and suggestions; Lauriane Fritsch, Boyan Bonev, and Giacomo Cavalli for SAM ESCs and SAM MEFs; Hervé Seitz, Adrien Décorsière, and Laure Garnier for critical reading of our draft manuscript; Chris Planque for advice; Philaé Gil and Pierre Nègre for technical assistance; Lionel Quentin and Nicolas Boucharel for performing mycoplasma tests; members of the IGF for continuous support. pB-CAGGS-dCas9-KRAB-MeCP2 was a gift from Alejandro Chavez & George Church (Addgene plasmid # 110824). HyperPiggyBac Transposase (pCMV-HA-HyperpBase) was a gift of Kosuke Yusa and Allan Bradley and was kindly provided by Thomas Dibling (Sanger, Cambridge, U.K). sgRNA(MS2) cloning backbone and Lenti sgRNA(MS2)_zeo backbone were a gift from Feng Zhang (Addgene plasmids # 61424 and 61427).

## AUTHOR CONTRIBUTIONS

T.B. conceived the project and designed the study with inputs from A.V.; I.C., M.C., and T.B. performed cell culture and qPCR experiments; A.L.D. and T.B. generated constructs; A.M. and. C.L. generated lentiviruses; C.H. generated DMH1-HCl; T.B. generated CRISPR cell lines; T.B. performed all of the analyses and drafted the manuscript; A.V. and T.B. revised the manuscript. All authors read and approved the final version of the manuscript.

## SUPPLEMENTAL FIGURE LEGENDS

**Supplemental Figure S1.**
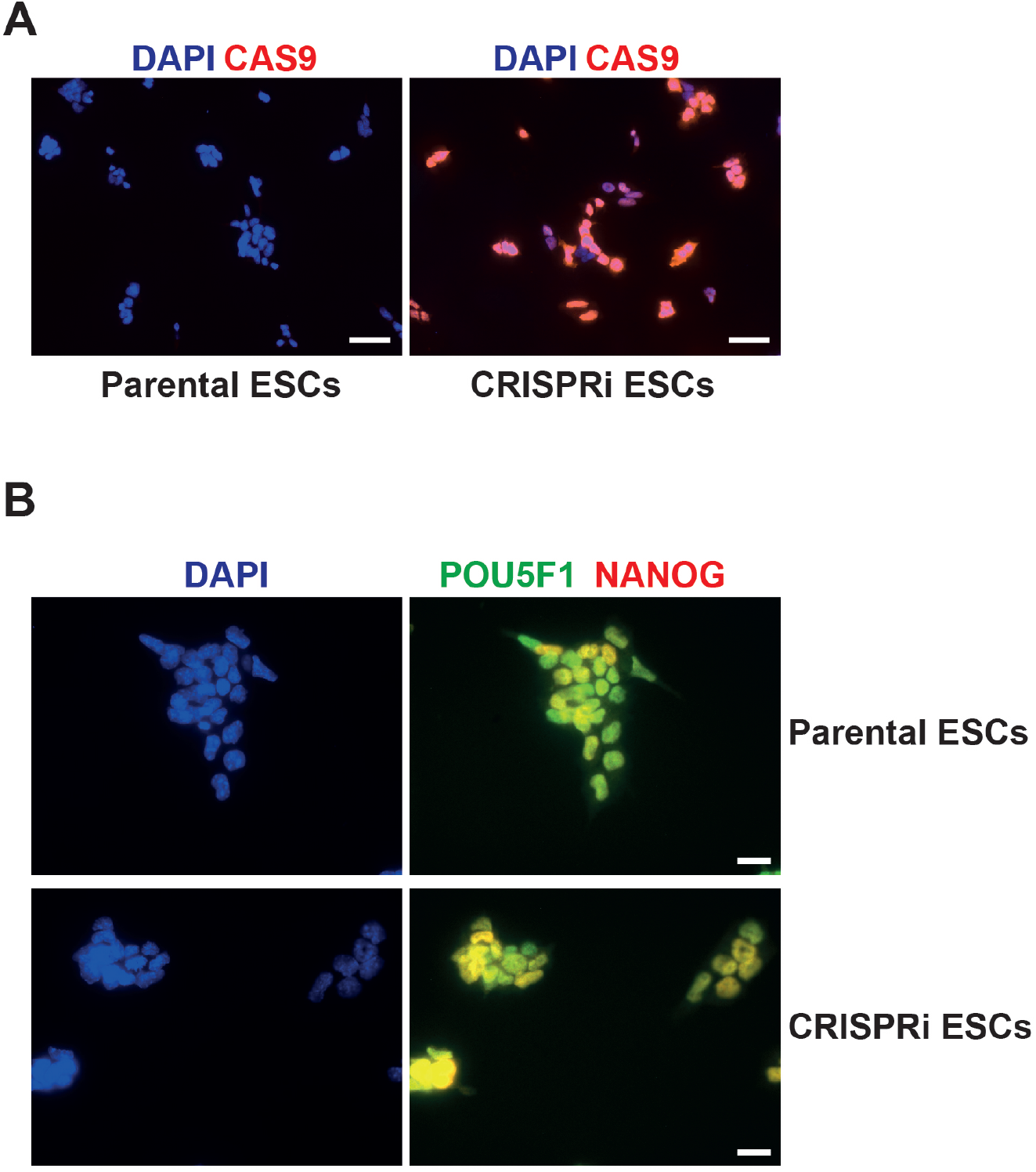
Characterization of the dCas9-KRAB-MeCP2 (CRISPRi) ESC line. (A) Expression of CAS9 in parental E14Tga2 ESCs and their CRISPRi derivatives. Scale bars: 50 µm. (B) Expression of pluripotency factors POU5F1 (green) and NANOG (red) in parental E14Tga2 ESCs and their CRISPRi derivatives. Scale bars: 20 µm.

**Supplemental Figure S2.**
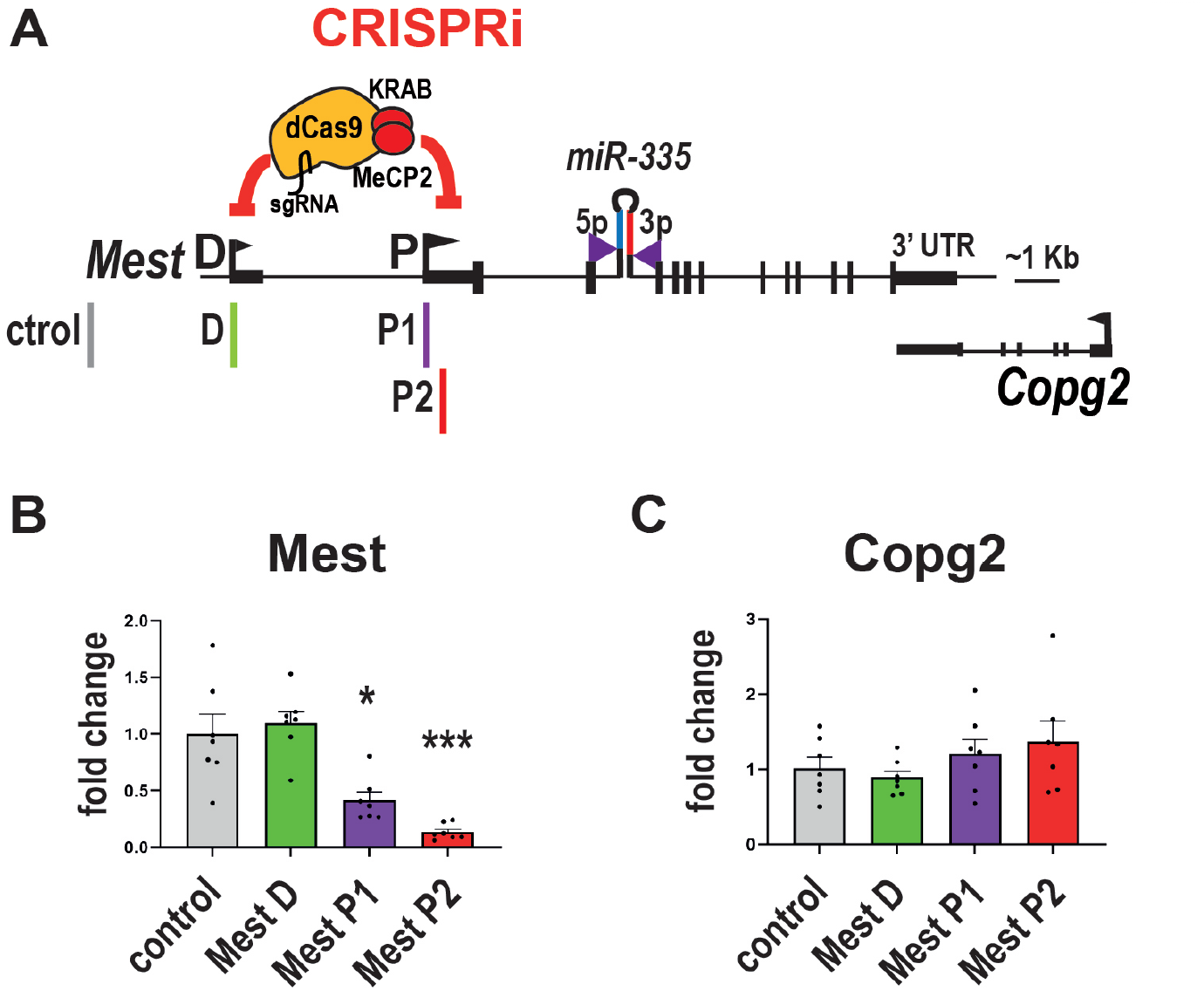
Efficient CRISPRi of *Mest* when targeting its proximal promoter. (A) Structure of mouse *Mest* gene with the CRISPRi module (dCa9-KRAB-MeCP2) directed to either the distal (*D*) (sgRNA D, green) or proximal (P) (sgRNAs P1 -purple- and P2 -red-) promoter of *Mest*. The sgRNA control (grey) has no match in the mouse genome. (B, C) Repression of *Mest* proximal promoter downregulates *Mest* expression (B) and does not affect neighboring *Copg2* expression (C). CRISPRi ESCs were transduced with lentiviruses expressing either sgRNA control, sgRNA D, sgRNA P1, or sgRNA P2. RNAs were measured by RT-qPCR. Data are mean ± sem of seven independent experiments and expressed as fold change over control sgRNA. *: p<0.05, ***: p<0.001 in Mann-Whitney test (comparison with sgRNA control values). Only the sgRNAs targeting the proximal promoter repressed *Mest*.

**Supplemental Figure S3.**
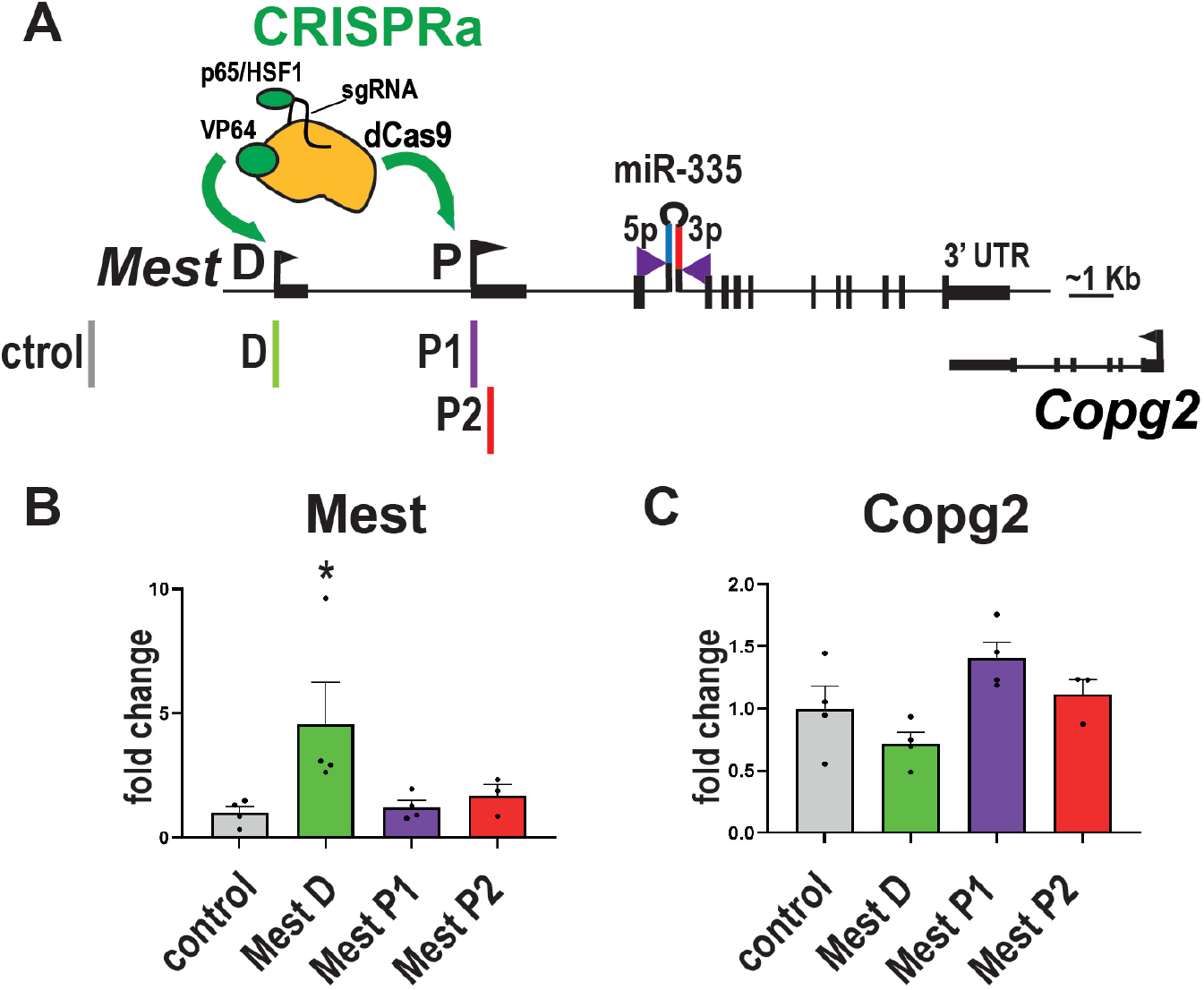
Efficient CRISPRa of *Mest* when targeting its distal promoter. (A) Structure of mouse *Mest* gene with the CRISPRa tool SAM targeting either the distal (sgRNA D, green) or the proximal (sgRNAs P1 -purple- and P2 -red-) promoter of *Mest*. The sgRNA control (grey) has no match in the mouse genome. (B, C) Transactivation of *Mest* distal promoter upregulates *Mest* expression (B) and does not affect neighboring *Copg2* expression (C). CRISPRa SAM ESCs were transduced with lentivirus expressing either SgRNA control, D, P1, or P2. *Mest* (B) and Copg2 (C) were measured by RT-qPCR. Data are mean ± sem of four independent experiments. *: p<0.1 in Mann-Whitney test. Only the sgRNA targeting the distal promoter upregulated *Mest* RNA.

**Supplemental Table S1.**
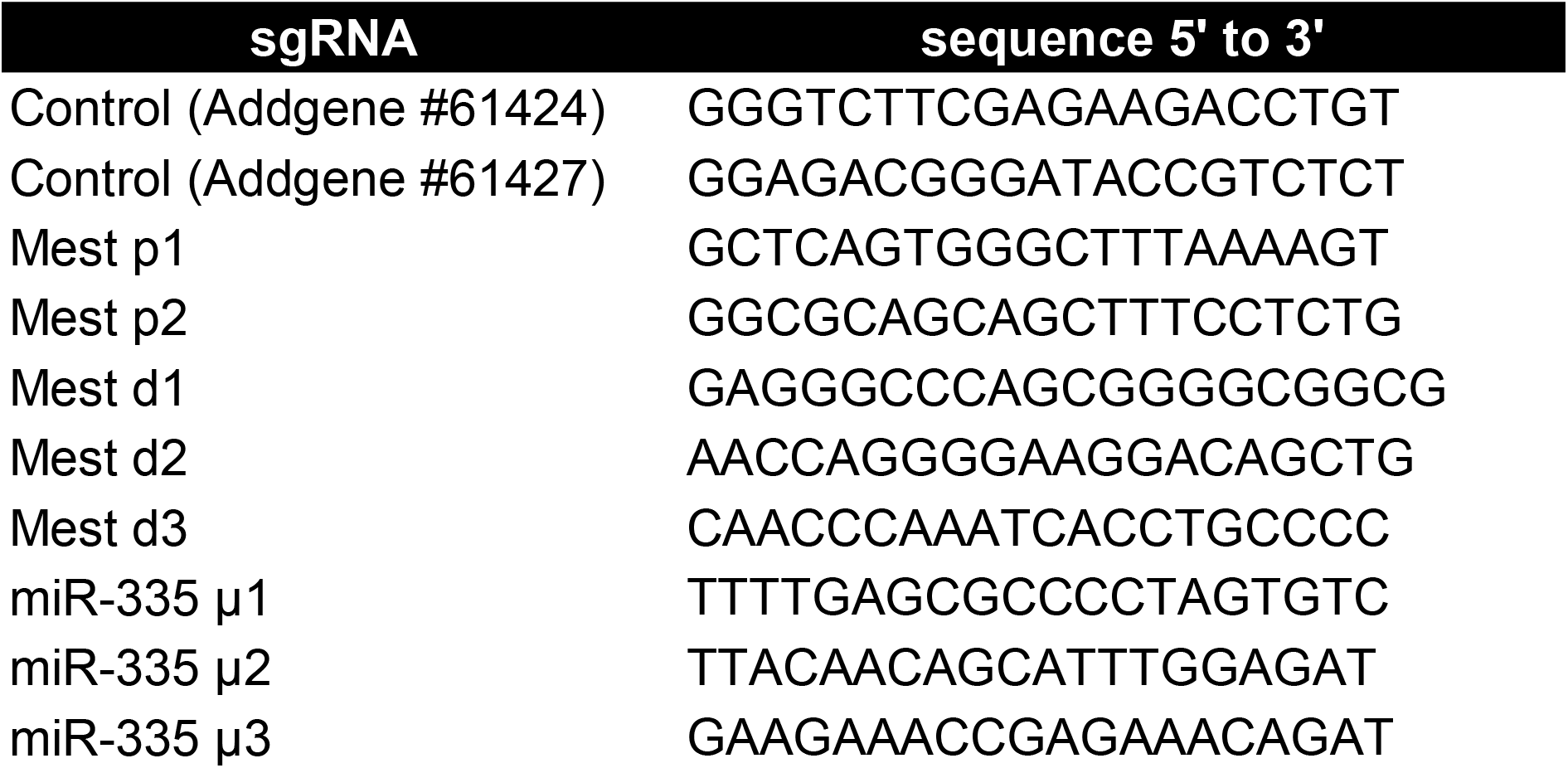
Sequences of sgRNAs.

**Supplemental Table S2.**
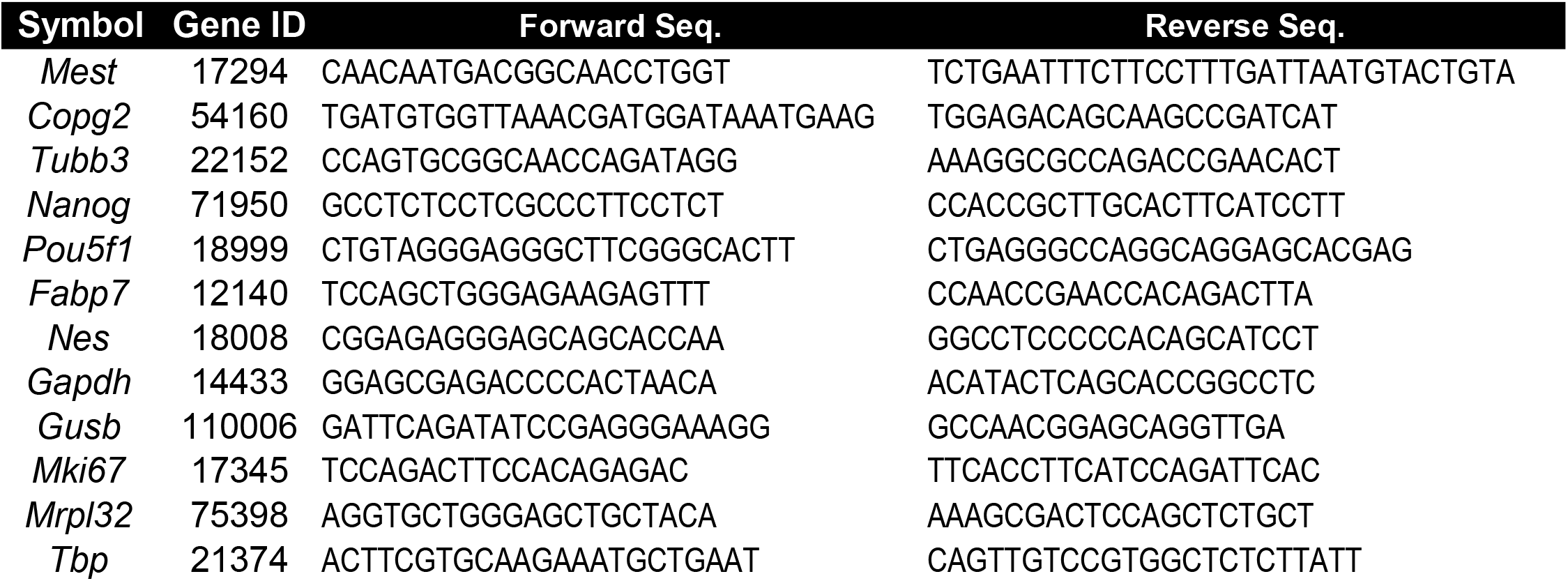
Primers used for qPCR assays.

